# A digital twin of freely moving mouse V1 learns a general visual code bridging classical physiology and active vision

**DOI:** 10.64898/2026.11.13.766486

**Authors:** Marius Schneider, Yuchen Hou, Cristopher M. Niell, Michael Beyeler

## Abstract

Decades of controlled experiments in passive, head-fixed animals have characterized the response properties of mouse primary visual cortex (V1) and their strong modulation by behavioral state. However, in natural vision the animal is active, continuously shaping its own visual input as it moves and coupling vision to behavior. To bridge this gap between classical controlled experiments and natural vision, we trained a multimodal digital twin to predict V1 spiking in freely moving mice from retinocentric video together with locomotion, pupil, and eye and head movements, then froze it and probed it *in silico* under controlled conditions. The representation it learned generalized across stimulus classes, animals, and recording regimes. Although never trained on parametric stimuli or given cell-type labels, the model recovered V1’s canonical tuning for orientation, spatial frequency and contrast, including the systematic differences between putative excitatory and inhibitory cells. A single model transferred to new mice from a few minutes of recording. Without retraining, it matched the representational geometry of an independent head-fixed dataset of natural movies as closely as two recorded mice match each other. Behavior modulated most V1 neurons, and clamping each variable in turn showed that locomotion and pupil dilation share a gain increase, while only locomotion shifts sensitivity toward high spatial frequencies. By learning this representation from the freely behaving animal and probing it under controlled conditions, freely moving digital twins bridge classical visual physiology and active vision.

## Introduction

Over the past decades, the response properties of mouse primary visual cortex (V1) have been characterized in remarkable detail using controlled parametric stimuli. These studies established canonical tuning properties such as orientation and direction selectivity, spatial-frequency tuning, contrast gain control, and their systematic differences across putative cell types (Niell and Stryker, 2008; Kerlin et al., 2010; Atallah et al., 2012; Busse et al., 2011). Subsequent work showed that these visual responses are strongly modulated by behavioral state, from locomotion and pupil dilation to spontaneous facial and body movements (Niell and Stryker, 2010; Ayaz et al., 2013; Aydin et al., 2018; Busse, 2018; Dadarlat and Stryker, 2017; Erisken et al., 2014; Horrocks et al., 2024; Neske et al., 2019; Saleem et al., 2013; Vinck et al., 2015; Stringer et al., 2019; Musall et al., 2019). Both the visual tuning and the behavioral modulation were measured almost entirely under head fixation, with the animal passively viewing experimenter-controlled stimuli.

Natural vision, however, is active, with the animal’s own movements continuously shaping the retinal input and embedding vision in a closed-loop action-perception cycle (Gibson, 1978; Skyberg and Niell, 2024; Niell et al., 2026). Head fixation removes head movement, and with it the eye-head coordination that produces natural gaze shifts. Advances in head-mounted cameras and motion tracking now make it possible to record these signals in freely moving mice, together with V1 activity, the visual scene, and locomotion (Meyer et al., 2018; Guitchounts et al., 2020; Parker et al., 2022). Such recordings have revealed that V1 activity depends on the animal’s eye and head position (Parker et al., 2022), that gaze shifts initiate a dynamic sequence of visual processing (Parker et al., 2023), and that these gaze shifts comprise distinct head-eye coordination motifs with motif-dependent V1 responses (Hou et al., 2026). But they make visual tuning hard to measure, because characterizing tuning requires the experimenter, not the animal, to determine what is seen.

Predictive modeling offers a way to regain this control. A digital twin, trained to reproduce the responses of a recorded neural population, can afterwards be probed in silico in place of the animal. These models are expressive deep networks, over-parameterized input-output mappings rather than mechanistic models of the circuit. Trained on large, curated sets of images or videos, such twins have been shown to generalize to stimuli outside their training distribution, and to synthesize the stimuli that most strongly drive individual neurons, verified by presenting those stimuli to the animal (Walker et al., 2019; Wang et al., 2025). So far, these models have been trained on head-fixed recordings. Xu et al. (2023) trained a digital twin on freely moving recordings, where behavior varies naturally and is richly sampled, with behavior as an explicit input alongside vision.

Building on that model, we trained a multimodal digital twin on recordings from freely moving mice (Parker et al., 2022), predicting V1 spiking from retinocentric video together with the animal’s eye and head movements, locomotion, and pupil diameter. A single core was trained jointly across animals, with a separate linear readout for each mouse’s neurons. We then froze the model and presented classical parametric stimuli while independently clamping each behavioral variable, characterizing both the visual and the behavioral tuning of its neurons. The framework thus combines the realism of natural behavior with the experimental control of classical physiology.

A shared core trained across animals learned a representation that transferred to held-out mice from a few minutes of recording. Probed with parametric stimuli, the model expressed canonical V1 tuning, including orientation and direction selectivity, spatial-frequency and contrast tuning, and their systematic differences across putative cell types. The same neurons were tuned to behavioral state. Locomotion and pupil diameter raised firing across most of V1, whereas head and eye position modulated firing with a preferred direction that differed from neuron to neuron. Clamping each variable in turn showed that locomotion and pupil dilation share a gain increase, while only locomotion redistributes sensitivity toward higher spatial frequencies. Finally, we replayed natural movies from the Allen Brain Observatory, recorded in head-fixed mice, through the frozen twin. The same representation, without retraining, matched the representational geometry of the real recordings as closely as two independently recorded mice matched each other, despite the different stimuli, recording setup, and animals. Together, these results show that a visual code learned from free exploration generalizes across stimulus classes, animals, and recording regimes.

## Results

### A shared visual-behavioral model predicts V1 activity in freely moving mice and transfers to new animals from little data

We trained a digital twin to predict the spiking of each V1 neuron from a freely moving mouse’s own world-view video, recorded with a head-mounted camera, together with its eye and head movements, locomotion, and pupil (Figure 1A,B). Building on Xu et al. (2023), who first trained such a model on these recordings, we asked whether a single representation could be shared across animals and capture V1 well enough to serve as a common model across freely moving mice.

**Figure 1:**
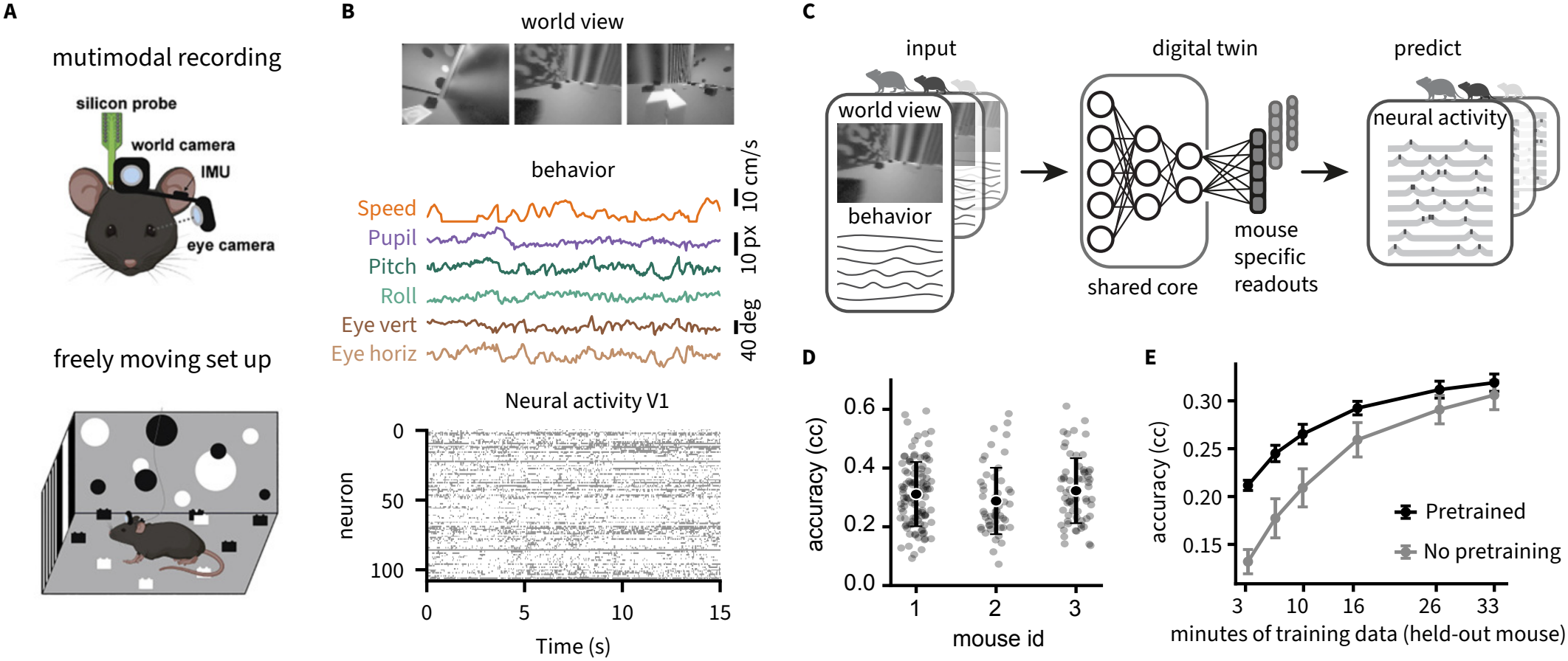
Predictive modeling framework for V1 activity in freely moving mice. **(A)** Multimodal recordings during freely moving behavior. Mice navigated an open arena while V1 activity was recorded with a silicon probe; retinocentric visual input was estimated from a head-mounted world camera, and behavioral variables (eye position, head pitch and roll, locomotion speed, pupil diameter) were tracked. **(B)** Example aligned data streams: world-view frames, behavioral traces, and the simultaneously recorded V1 population raster. **(C)** Predictive model architecture. World-view video and behavioral variables are processed by a shared visual– behavioral core, and a mouse-specific linear readout maps the shared representation onto recorded neurons to predict activity. **(D)** Single-neuron prediction accuracy across neurons and mice, quantified as the cross-correlation (cc) between predicted and recorded activity. **(E)** Cross-mouse generalization through pretraining. The shared core was pretrained on a subset of mice and adapted to a held-out mouse by fine-tuning while training a new readout; pretraining improved prediction over training from scratch, particularly in the low-data regime.

The architecture comprised a shared visual-behavioral core trained jointly across animals and mouse-specific linear readouts mapping the shared representation onto recorded neurons (Figure 1C). This design enforces a common representational space across mice while allowing differences in recorded neural populations to be captured at the output stage. The worldview video was warped frame by frame according to the measured eye and head position, so the visual input reaching the core followed the animal’s gaze and was effectively retinocentric (Methods). Across neurons and mice, the model predicted single-neuron responses at a mean cross-correlation (cc) of 0.31 between predicted and recorded spike counts (223 neurons from three mice; Figure 1D). Evaluated on 2s-smoothed responses, the mean correlation was 0.59, matching the previous digital twin trained on these recordings (Xu et al., 2023); all other correlations in this paper are computed on unsmoothed responses.

We next asked whether pretraining the shared core yields a data-efficient initialization for new animals. In a leave-one-mouse-out design, the core was pretrained on a subset of mice and then adapted to a held-out mouse by fine-tuning model parameters while training a new readout. Pretraining improved prediction performance relative to training the same architecture from scratch on the held-out mouse, with the largest gains in the low-data regime (Figure 1E). With 3.3 minutes of recording from a new mouse, the pretrained model reached cc = 0.21, 60% higher than training the same architecture from scratch on the same data (cc = 0.13), and the two converged only with substantially more data (0.32 versus 0.31 at 33 minutes). This indicates that pretraining captures structure conserved across individuals.

Finally, we tested whether the behavioral inputs contribute beyond vision alone by comparing the full multimodal model with a vision-only control (Figure S1). Including behavioral variables improved single-neuron prediction for 169 of 223 neurons (76%; mean cc = 0.309 versus 0.300, paired Wilcoxon *p* = 4.0 × 10^−18^; Figure S1), confirming that behavior carries information about V1 activity that vision alone does not capture. All subsequent analyses therefore use the full visual-behavioral model. The shared model thus captures V1 responses during free behavior, and the representation it learns transfers to a new animal from a few minutes of recording.

### Canonical, cell-type-specific V1 tuning is recovered from freely moving visuomotor experience

We classified the recorded neurons as broad- or narrow-waveform (putative excitatory and inhibitory; Figure 2A) from the shape of their spike waveforms (Niell and Stryker, 2008; Barthó et al., 2004). After training on the freely moving data, we froze the model’s parameters and replayed drifting gratings through it, applying classical tuning analysis to ask whether the canonical, cell-type-specific tuning established by decades of V1 physiology was present in a representation learned from free behavior. The model was never trained on the controlled parametric stimuli used to characterize tuning, nor given cell-type labels.

**Figure 2:**
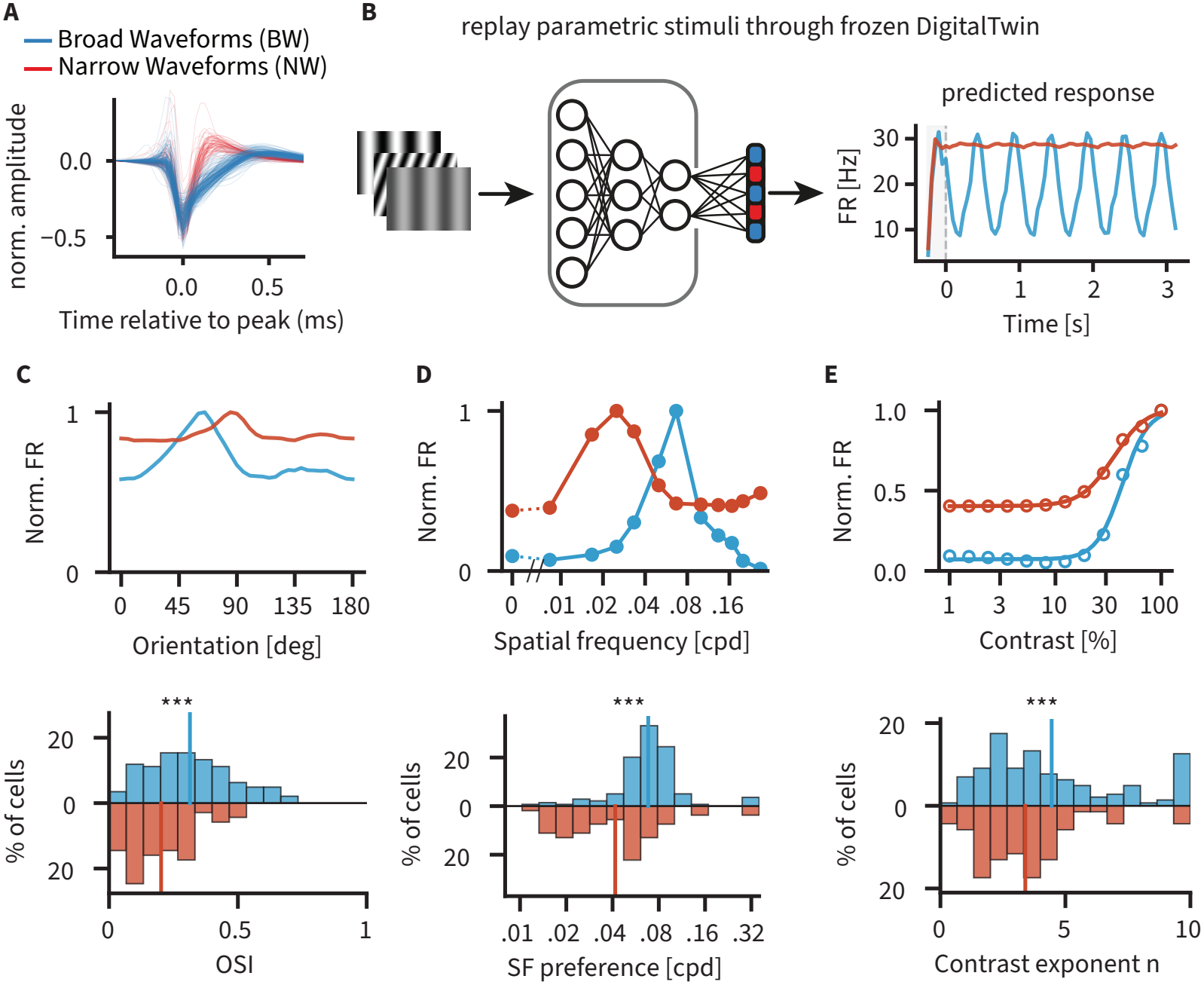
Canonical, cell-type–specific V1 tuning properties assessed by *in silico* probing. **(A)** Normalized extracellular spike waveforms for broad-waveform (putative excitatory) and narrow-waveform (putative inhibitory) neurons in V1. **(B)** Schematic of the *in silico* replay protocol: after training on freely moving behavior, model parameters were frozen and the model was probed with drifting grating stimuli while behavioral inputs were held constant. Right, predicted firing rates of two representative broad- and narrow-waveform units in response to drifting gratings. **(C)** Orientation tuning. Top, example orientation tuning curves for representative broad- and narrow-waveform units. Bottom, population distributions of the orientation selectivity index (OSI). **(D)** Spatial-frequency tuning. Top, example spatial-frequency tuning curves. Bottom, population distributions of preferred spatial frequency, band-pass units only. **(E)** Contrast responses. Top, normalized contrast–response functions of example neurons. Bottom, population distributions of the fitted contrast–response exponent. Distributions in (C)-(E) are mirrored histograms for the two cell types with the group mean marked; group comparisons use a permutation test on the difference of means. Direction selectivity and spatial-frequency bandwidth are shown in Figure S2.

Figure 2B (right) shows representative predicted firing-rate traces during drifting gratings, with strong phase modulation in the broad-waveform example and higher baseline firing with weaker phase locking in the narrow-waveform example. Example orientation tuning curves demonstrate sharper selectivity in the representative broad-waveform unit and weaker selectivity in the narrow-waveform unit (Figure 2C, top). At the population level, orientation selectivity indices (OSI) fell within established physiological ranges, and broad-waveform units exhibited significantly higher OSI than narrow-waveform units (mean 0.31 versus 0.20; permutation test, *p* = 5 × 10^−5^; Figure 2C, bottom), in agreement with measurements in head-fixed mouse V1 (Niell and Stryker, 2008; Kerlin et al., 2010; Runyan et al., 2010). Direction selectivity showed a similar pattern. Broad-waveform units exhibited higher direction selectivity than narrow-waveform units (mean DSI 0.13 versus 0.09; *p* = 9 × 10^−4^; Figure S2A), consistent with reported differences between excitatory and inhibitory populations in mouse V1 (Niell and Stryker, 2008).

Spatial-frequency tuning included both band-pass and lowpass profiles, with the majority of units exhibiting band-pass tuning; low-pass profiles were about eight times as common among narrow-waveform units (21.7%, 15*/*69) as among broad-waveform units (2.8%, 4*/*143; Figure S2B). Among band-pass units, narrow-waveform units displayed broader spatial-frequency tuning than broad-waveform units (mean bandwidth 3.43 versus 2.82 octaves; *p* = 0.003; Figure S2B) and preferred lower spatial frequencies (geometric mean 0.042 versus 0.068 cycles per degree; *p* = 5 × 10^−5^; Figure 2D), in agreement with head-fixed *in vivo* measurements in mouse V1 (Niell and Stryker, 2008; Andermann et al., 2011).

Finally, neurons exhibited saturating contrast-response functions across the tested contrast range (Figure 2E, top). Contrast response exponents spanned a broad range and were larger in broad-waveform than in narrow-waveform units (mean 4.45 versus 3.40; *p* = 0.007; Figure 2E, bottom), as reported for putative excitatory and inhibitory neurons in mouse V1 (Atallah et al., 2012).

These controlled *in silico* experiments show that a representation learned from freely moving visuomotor experience expresses the canonical, cell-type-specific tuning of mouse V1, consistent with decades of head-fixed visual physiology, and that it does so for a stimulus class the model never encountered during training. V1 activity is also shaped by behavior, from locomotion and pupil dilation to the animal’s eye and head movements (Niell and Stryker, 2010; Vinck et al., 2015; Parker et al., 2022). Because the model jointly represents vision and behavior, we next mapped the behavioral tuning of these same neurons across all six variables, isolating the contribution of each while holding the others fixed, a separation that is difficult in freely moving animals, where these variables naturally co-vary.

### Behavioral state modulates most V1 neurons through two distinct regimes

Holding the visual input fixed, we swept each behavioral variable across its natural range while clamping the others, and read out how it changed the predicted firing of every neuron (Figure 3A). A single modulation index captured each neuron’s response to each variable, positive when higher values drove firing up and negative when they suppressed it. Across all six variables, most neurons were significantly modulated by behavior (82–92% of neurons; shape-aware permutation test; Figure S3A), so the twin predicts that behavior modulates most V1 neurons. The form of that modulation, however, separated the variables into two regimes (Figure 3B,C).

**Figure 3:**
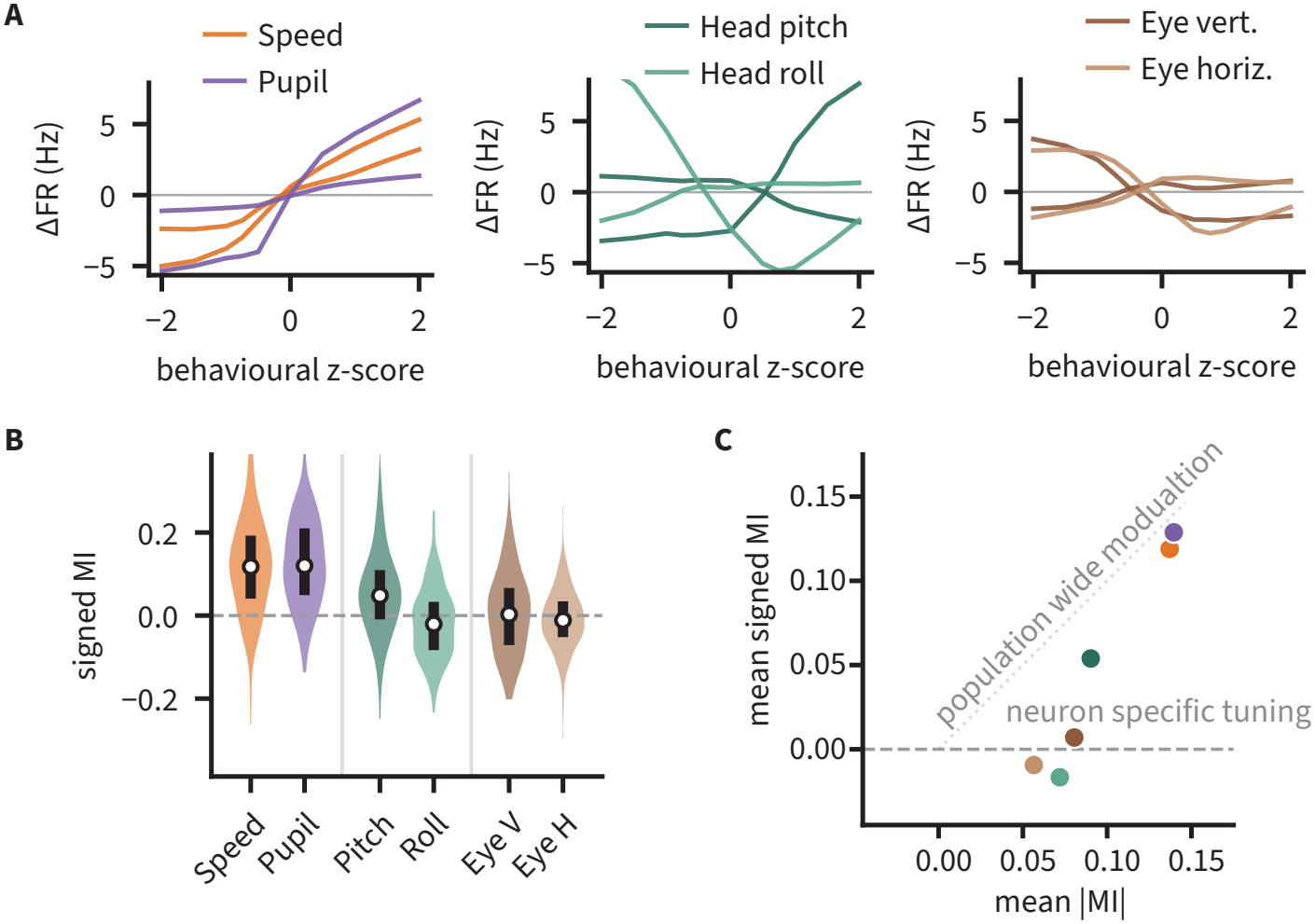
Behavioral tuning of V1 neurons in the digital twin. **(A)** Example tuning curves. For each of six behavioral variables, grouped as arousal/state (locomotion speed, pupil diameter), head position (pitch, roll), and eye position (vertical, horizontal), the change in predicted firing rate (ΔFR) of two example neurons as a function of the behavioral variable (z-scored across its range). For the head- and eye-position variables the two examples were selected to have opposite signed modulation, illustrating that the preferred direction differs across neurons. All panels share axes. **(B)** Population distributions. Distribution of the per-neuron modulation index (MI) for each variable across all *N* = 212 neurons (white dot, median; black bar, interquartile range). The number above each violin gives the percentage of significantly modulated neurons with positive MI. Locomotion speed and pupil diameter shift toward positive values, whereas the head- and eye-position variables are centered on zero. **(C)** Strength versus direction. Mean signed MI against mean |MI |across neurons, one point per variable. Points on the diagonal indicate population wide modulation, where every neuron is modulated in the same direction; points on the horizontal indicate a neuron-specific preference, with as many neurons modulated positively as negatively. The arousal variables lie near the diagonal, whereas head roll and the two eye-position variables fall to the horizontal at comparable |MI|; head pitch is intermediate.

Locomotion speed and pupil diameter acted as unipolar arousal signals. Neurons fired more at higher values of each, and nearly all were modulated in the same direction (median MI = 0.12 for both; 88% and 92% of significantly modulated neurons positively modulated; *N* = 212, Wilcoxon signed-rank *p* = 5.2 × 10^−27^ and *p* = 1.2 × 10^−30^). Head and eye positions instead modulated firing around a neutral posture, consistent with the tuning of freely moving V1 neurons for eye and head position (Parker et al., 2022). Individual neurons increased or decreased their rate depending on the direction of the deviation, but the preferred direction differed from neuron to neuron, so the signed index averaged to near zero (mean MI between − 0.017 and +0.007 for head roll and the two eye variables) even though these variables significantly modulated just as many neurons (82–91%). Head pitch was intermediate, combining a monotonic component with deviation-dependent tuning. Plotting each variable’s mean signed MI against its mean |MI| separates the two regimes directly (Figure 3C), with speed and pupil near the identity line and head roll and both eyeposition variables on the horizontal at comparable |MI|. The cell-type structure recovered in the visual tuning extended to behavioral modulation. Narrow-waveform (putative inhibitory) neurons were more strongly modulated by locomotion speed than broad-waveform neurons (median MI 0.17 versus 0.11; Mann–Whitney U, *p* = 4.2 × 10^−4^), an effect specific to running speed among the six variables (Figure S3B), consistent with previous observations in head-fixed recordings of mouse V1 (Niell and Stryker, 2010; Polack et al., 2013).

Locomotion and pupil dilation are both associated with elevated V1 activity and co-vary during natural behavior (Niell and Stryker, 2010; Vinck et al., 2015). We next asked how each reshapes V1’s visual tuning, beyond the overall gain they share.

### Locomotion and pupil dilation increase V1 gain, and locomotion redistributes spatial-frequency sensitivity

Behavioral state could simply scale visual responses, or it could reshape which visual features V1 represents. We separated these possibilities by replaying identical drifting gratings through the frozen model while clamping the behavioral inputs, so that any change in the response reflected behavioral state rather than the visual stimulus (Figure 4A), with every comparison made against a common baseline in which locomotion was stationary and the remaining variables sat at their mean. Example neurons show both outcomes. Orientation tuning was unchanged across states (Figure 4C), while spatial-frequency responses shifted toward higher frequencies during locomotion (Figure 4E).

**Figure 4:**
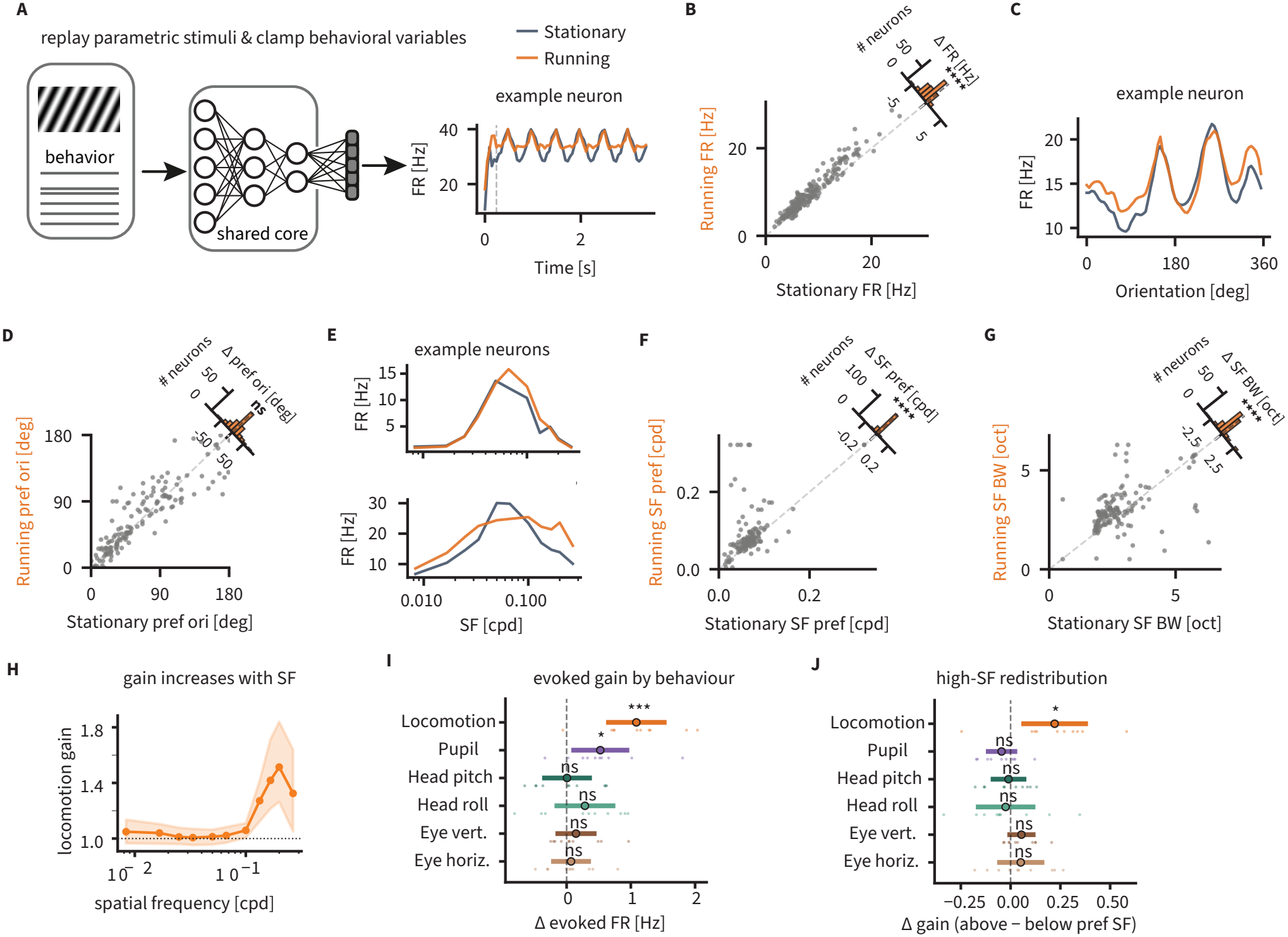
Locomotion and pupil dilation increase V1 gain, and locomotion redistributes spatial-frequency sensitivity. **(A)** Behavioral inputs were clamped while identical drifting gratings were replayed through the frozen model, isolating the effect of state from the visual stimulus. In (B)–(H) locomotion was clamped to stationary (0 cm/s) or running (4 cm/s) with the other five variables held at their mean. Right, predicted firing of an example neuron across the two states. **(B)** Visually evoked firing rate of each neuron in stationary versus running states, with the distribution of the change (ΔFR); locomotion increases evoked firing across the population. **(C)** Orientation tuning of an example neuron, unchanged between stationary and running states. **(D)** Preferred orientation in stationary versus running states across neurons, with the distribution of the change (Δ preferred orientation, not significant). **(E)** Spatial-frequency tuning of two example neurons. **(F)** Preferred spatial frequency in stationary versus running states, with the distribution of the change; locomotion shifts preferred spatial frequency upward. **(G)** Spatial-frequency bandwidth in stationary versus running states, with the distribution of the change; locomotion broadens SF tuning. **(H)** Locomotion gain as a function of stimulus spatial frequency; median across band-pass neurons (*n* = 145). **(I)** Specificity of the gain increase. The same manipulation was applied to each of the six behavioral variables in turn, every one swept the same distance in z-scored units, and the whole analysis was repeated in ten digital twins retrained from scratch with different random seeds. Change in evoked firing rate in each network (points), with the mean across networks and its 95% confidence interval. **(J)** Specificity of the high-spatial-frequency redistribution. Change in high-spatial-frequency bias, defined for each neuron as the gain above its preferred spatial frequency minus the gain below it, across the same six variables, plotted as in (I). Asterisks denote ^*\**^ *p <* 0.05, ^*\*\**^ *p <* 0.01, ^*\*\*\**^ *p <* 0.001, ^*\*\*\*\**^ *p <* 10^−4^; ns, not significant. In (B), (D), (F) and (G) they refer to Wilcoxon signed-rank tests across neurons. In (I) and (J), where each point is one retrained network, they refer to one-sample *t*-tests of the ten network values against zero.

Both locomotion and pupil dilation increased visually evoked firing rates across the population (locomotion +1.15 Hz, *n* = 212, *p <* 10^−29^; Figure 4B; pupil in Figure 4I), in line with the known enhancement of V1 responses by locomotion and arousal (Niell and Stryker, 2010; Vinck et al., 2015). This gain left orientation tuning intact. Preferred orientation was unchanged across states (80% of orientation-tuned neurons within 30^°^, *n* = 161; Figure 4D), so behavioral state scaled responses without reorganizing orientation preference.

Locomotion reshaped visual processing beyond gain. Its gain was not uniform across stimulus spatial frequency. Computed within each neuron, it was near unity at the lowest spatial frequency tested (1.09 at 0.008 cycles per degree) and rose to a peak of 1.53 at 0.233 cycles per degree (*n* = 145 band-pass neurons, *p* = 1.9 × 10^−9^; Figure 4H). Example neurons show the same shift toward finer detail (Figure 4E). Locomotion therefore shifted preferred spatial frequency upward (median 0.064 to 0.073 cycles per degree; Figure 4F) and broadened spatial-frequency tuning (median 2.45 to 2.84 octaves; Figure 4G), redistributing sensitivity toward finer spatial detail rather than scaling all spatial frequencies equally. This is consistent in direction with the finding that locomotion improves spatial resolution in mouse V1 (Mineault et al., 2016), although there the effect was a difference across the neuronal population rather than the within-neuron reweighting seen here (see Discussion).

This redistribution was specific to locomotion. To test this we swept each of the six behavioral variables in turn by the same distance, 1.5 standard deviations from its own tenth percentile. Any single trained network is one of many that fit these recordings equally well, so an effect it shows need not follow from the neural data itself. We therefore repeated the whole analysis in ten digital twins retrained from scratch with different random seeds, and counted an effect as real only if it held across them. As a further control, three models trained with the locomotion trace shuffled in time showed neither effect (Methods).

Only locomotion and pupil dilation raised evoked firing rate reliably (mean across networks +1.09 and +0.53 Hz; *p* = 3 × 10^−4^ and *p* = 0.02), while the postural and eye-position variables did not (Figure 4I). The two variables that share this gain increase separate on spatial frequency. Locomotion was the only variable whose high-spatial-frequency bias differed from zero across networks (mean +0.22, *p* = 0.01), shifting in the same direction in nine of the ten, whereas pupil dilation showed no consistent shift (Figure 4J).

Locomotion and pupil dilation both increase responsiveness, but only locomotion redistributes sensitivity toward finer spatial detail. This separates a general arousal-related gain from a locomotion-specific change in which spatial features V1 represents most strongly.

### A representation learned from free exploration generalizes to head-fixed natural-movie viewing

Replaying drifting gratings through the frozen twin showed that it expressed V1’s canonical tuning, including the well-characterized differences between cell types, for a stimulus class it was never trained on (Figure 2). The natural movies of the Allen Brain Observatory provide a harder test, because they were shown to head-fixed mice viewing passively, so the stimuli, the behavioral state, the recording setup and the animals all differ from anything the twin saw in training.

We replayed these movies through the frozen twin, with its behavioral inputs set to zero, and asked whether the V1 population it predicted was organized like the real one (Figure 5A,B). Rather than compare neuron by neuron, we compared the geometry of the two populations, how similarly each represents the same moments of the movie, and asked how close the twin came to real V1 relative to how close two real mice come to each other (Figure 5C). Because measurement noise attenuates the model-to-animal and animal-to-animal comparisons by different amounts, we corrected for it before comparing them (Methods).

**Figure 5:**
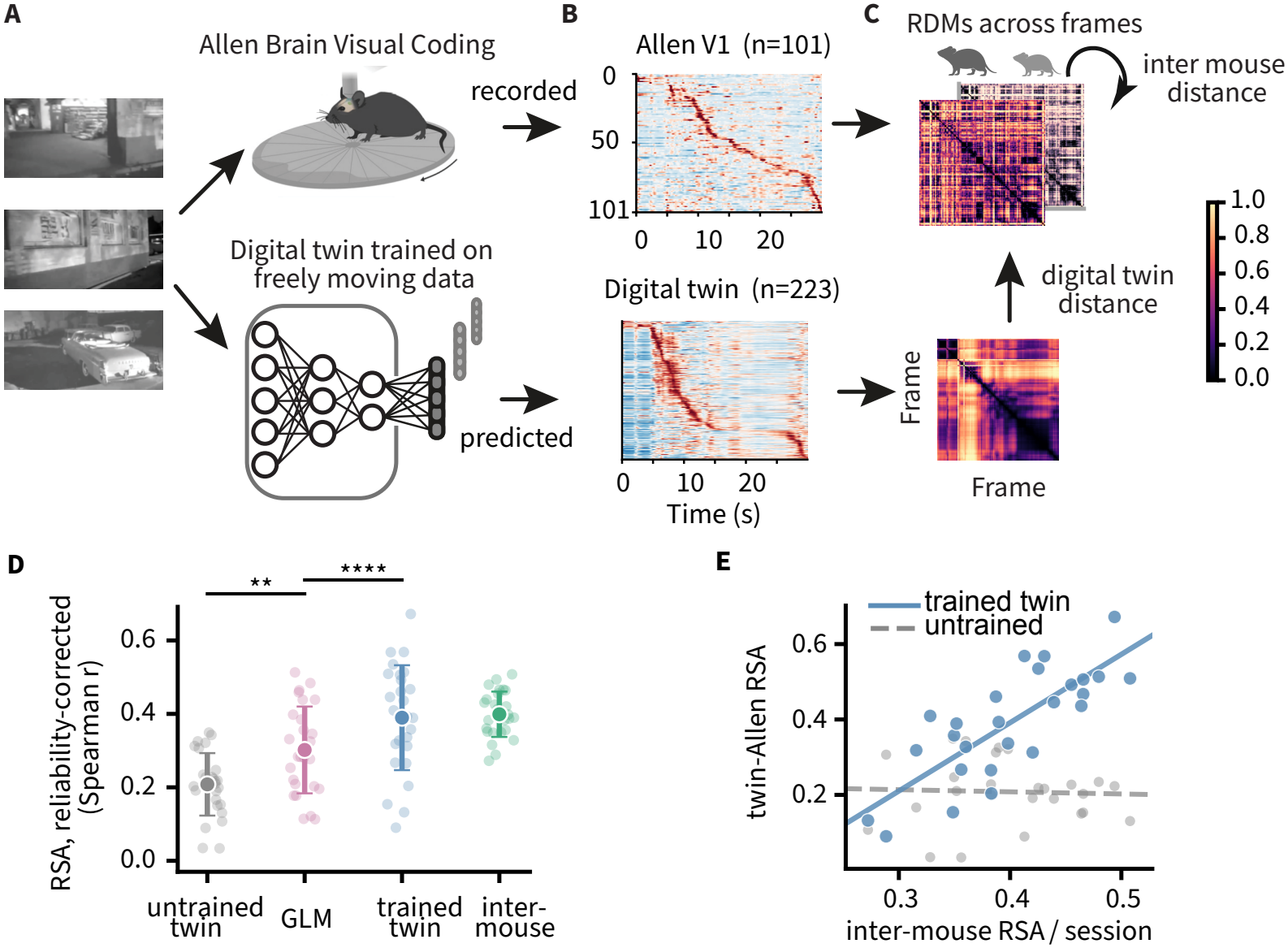
Generalization to an independent head-fixed natural-movie dataset. **(A)** Natural movies from the Allen Brain Observatory Visual Coding dataset were presented to head-fixed mice (recorded) and replayed through the digital twin trained on freely moving data (predicted). **(B)** Example population responses to a natural movie: recorded Allen V1 neurons (*n* = 101) and predicted digital-twin neurons (*n* = 223), each sorted by time of peak response. **(C)** Representational dissimilarity matrices (RDMs) across movie frames, computed from the predicted twin responses and from recorded Allen responses; the twin-to-Allen distance is compared with the distance between independently recorded Allen mice. **(D)** Representational similarity (Spearman correlation between RDMs), corrected for RDM unreliability. The trained twin (*ρ* = 0.39) exceeds both a randomly initialized model (0.21) and a GLM trained on the same freely moving data (0.30; per-session paired test, *p* = 6.2 × 10^−6^, twin higher in 23 of 26 sessions), and reaches the agreement between independently recorded mice (0.40), sitting at the 48th percentile of the mouse-to-mouse distribution (two one-sided tests against a margin set by the spread between mice, *p* = 3.8 × 10^−5^). **(E)** Across recording sessions, the twin-to-Allen similarity tracks the inter-mouse similarity (Spearman *ρ* = 0.82, *p* = 3.4 × 10^−7^), so the twin agrees with Allen V1 where Allen mice agree with one another; an untrained model does not.

The twin’s predicted responses matched the representational geometry of the recorded Allen data as closely as two independently recorded mice matched each other (*ρ* = 0.39 versus 0.40; equivalence test, *p* = 3.8 × 10^−5^). A randomly initialized model reached 0.21 and a GLM encoding model trained on the same freely moving data reached 0.30, so the GLM recovered about half of the twin’s advantage over an untrained network (Figure 5D). Session by session, the twin agreed with Allen V1 precisely where independently recorded mice agreed with one another (*ρ* = 0.82; Figure 5E), whereas an untrained model did not.

The representation learned from free exploration therefore generalizes beyond the stimuli it was trained on and beyond the regime in which it was learned, matching V1 recorded in different animals, in a different laboratory, under passive head-fixed viewing.

## Discussion

Head-fixed experiments deliver the stimulus control that defines V1’s canonical tuning, but at the cost of natural behavior; freely moving recordings restore natural, active vision, but give up that control. We combined both by training a digital twin to predict V1 activity from vision and behavior during free exploration, then probing its learned representations *in silico* under controlled conditions. The twin recovered the canonical, cell-type-specific tuning of mouse V1, and the representation it learned generalized beyond the animals and setup it was trained on, matching an independent head-fixed dataset as closely as two mice agree. Using this validated, general model, we then asked how behavioral state reshapes visual processing, finding that locomotion and pupil dilation share a gain increase while only locomotion enhances high-spatial-frequency sensitivity. By learning this representation from the freely behaving animal and probing it under controlled conditions, the twin bridges classical visual physiology and active vision.

### Canonical V1 tuning is recovered from a model trained only on freely moving data

The twin reproduced V1’s canonical, cell-type-specific tuning, including its excitatory–inhibitory differences, although it was never trained on the controlled parametric stimuli used to characterize tuning, nor given a tuning or cell-type label. Broad- and narrow-waveform neurons were separated only after training, from their recorded spike waveforms, yet they reproduce the published ordering on orientation selectivity, direction selectivity, spatial-frequency tuning and contrast. Recovering this tuning from freely moving data is not trivial, because the animal controls its own input and the repeated stimuli of classical measurement are unavailable; the twin instead learns a model that can afterwards be probed *in silico* with any stimulus. Canonical tuning therefore becomes measurable in recordings where no parametric stimuli were ever presented, and in preparations where head fixation is not an option.

### The learned representation generalizes across animals and recording regimes

The twin’s learned representation generalized far beyond its training data. The Allen Brain Observatory movies were recorded from different, head-fixed mice viewing passively, and their image statistics are unlike the self-generated arena views the twin trained on. Yet when we fed these movies through the frozen twin, its predicted V1 population matched the recorded Allen data as closely as two independently recorded mice matched each other.

This generalization was obtained without the large, high-diversity stimulus sets used to train head-fixed foundation models of visual cortex (Wang et al., 2025); the visual input generated by an animal exploring a single small arena was sufficient. As the mouse moves, it views each scene feature from continuously changing angles and distances, so active sampling generates its own visual diversity. Trained on freely moving data, such twins also inherit both visual and behavioral structure, which makes them a tool for studying how behavior reshapes visual representations. Whether active, self-generated sampling provides a more efficient training signal than curated stimulation, or the two are simply complementary, cannot be settled here and would require training matched models under each regime on the same recordings.

### Behavioral state raises gain broadly but reshapes spatial-frequency tuning only during locomotion

Locomotion and pupil dilation both increased visually evoked firing in V1, in line with long-standing work on locomotion- and arousal-related gain (Niell and Stryker, 2010). These two variables co-vary during natural behavior; earlier work separated their contributions by exploiting periods when they diverge, such as changes in arousal during quiet wake-fulness, and found that locomotion and arousal make distinct contributions to cortical state and visual encoding (Vinck et al., 2015). Clamping each variable independently in the model separated them directly and showed that the gain increase is shared by both arousal-related variables rather than specific to locomotion. When every variable was displaced by the same amount, only locomotion and pupil dilation raised firing, while head pitch, head roll and eye position did not.

What distinguished locomotion was not this overall gain but where in the spatial-frequency spectrum it acted. During locomotion, gain was largest at high spatial frequencies, enhancing responses to fine spatial detail, shifting preferred spatial frequency upward and broadening tuning, whereas pupil dilation raised gain by a comparable amount without this dependence. Gain and spatial-frequency reweighting are thus separable. A behavioral state can amplify V1 responses without changing which spatial features are most strongly represented, and locomotion is set apart by coupling the two. This redistribution is a prediction of the model and remains to be tested in vivo.

This locomotion-specific redistribution relates to, but is not identical with, earlier observations. Mineault et al. (2016) found that locomotion gain is larger for neurons tuned to higher spatial frequencies, a difference across the neuronal population. In our model the redistribution instead arises within individual tuning curves, with gain rising with stimulus spatial frequency in each neuron, so the effect reflects a within-neuron reweighting rather than only a sorting of gain across neurons by preferred spatial frequency.

Why locomotion specifically would enhance high-spatial-frequency responses remains unclear. One possibility is compensatory. Self-motion raises retinal image speed, and temporal integration in early vision attenuates high spatial frequencies more under fast motion, so greater high-spatial-frequency sensitivity during locomotion could partly offset this loss. This interpretation is speculative, and the redistribution might instead reflect a greater demand for fine spatial detail during active movement, or some combination of the two.

### Limitations and outlook

Three limitations qualify these results. First, the behavioral manipulations are *in silico* clamps, predictions of how V1 would respond rather than direct measurements, and will ultimately need testing against closed-loop experiments. The locomotion-specific spatial-frequency redistribution reproduces in nine of ten networks retrained from scratch (Figure 4I,J), which shows that it follows from the recordings rather than from one training run, but it remains untested *in vivo*. Second, the freely moving recordings come from a single dataset of three mice, though the generalization to an independent Allen dataset argues that the learned representation is not idiosyncratic to them. Third, although the twin generalizes well from recordings made in a single small arena, we cannot yet attribute this to active, self-generated sampling itself, because the freely moving and head-fixed regimes differ in several respects at once.

More broadly, freely moving digital twins make natural behavior accessible to the controlled experiments of classical physiology. Because the model captures vision and behavior jointly, it can ask how each behavioral variable shapes visual processing while holding the others fixed. And because a single representation generalizes across mice and transfers to an independent head-fixed dataset, the framework can be reused across animals and recording conditions rather than rebuilt for each. As recordings that pair neural activity with behavior become more widespread, freely moving digital twins offer a general way to study how the brain combines vision and behavior during active, real-world vision, in V1 and beyond.

## Acknowledgements

This work was supported by the National Institute of Neurological Disorders and Stroke of the National Institutes of Health under Award Number R01-NS121919. The content is solely the responsibility of the authors and does not necessarily represent the official views of the National Institutes of Health.

## Methods

### Experimental data

We used extracellular recordings from mouse primary visual cortex (V1) during free behavior, originally described in Parker et al. (2022). Briefly, 64- or 128-channel linear silicon probes (Diagnostic Biochips P64-3 or P128-6) mounted in custom 3D-printed drives were implanted in the left V1 of C57BL/6J mice. Probe placement was guided by widefield imaging to target the approximate center of left monocular V1. During recording sessions, mice first underwent head-fixed visual stimulation (white-noise receptive-field mapping) and were then transferred to a visually complex arena (~48 × 37 × 30 cm) containing black-and-white blocks, oriented gratings on the walls, and a monitor displaying moving spots.

Throughout free movement, six behavioral variables were continuously tracked: (1) locomotion speed, derived from a top-down tracking camera; (2–3) head pitch and roll, measured by a head-mounted inertial measurement unit (IMU); (4) pupil diameter (eyerad), extracted via DeepLabCut-based tracking of eight points around the pupil from a miniature eye camera; and (5–6) horizontal (*θ*) and vertical (*ϕ*) eye position, computed from the same eye-camera pupil fits. A forward-facing world camera (~120^°^ field of view) simultaneously captured the visual scene from the mouse’s perspective. All data streams were temporally aligned to 48 ms bins (~20.8 Hz) using system timestamps acquired in Bonsai (Lopes et al., 2015).

We used data from three recording sessions (mouse IDs: 070921_J553RT, 110421_J569LT, 101521_J559NC). Neural spike counts were binned at the common 48 ms frame rate and served as the prediction target. Data for each mouse were split into training (70%) and test (30%) sets, with each split further divided into 10 segments for efficient data loading. Cell-type classification (broad-waveform putative excitatory vs. narrow-waveform putative inhibitory) and recording depth were obtained from the original dataset metadata (Parker et al., 2022).

### Behavioral preprocessing

Missing values (NaN frames) in behavioral variables, which arise mainly when pupil tracking fails, were replaced by zeros. Zeros in the eye variables therefore denote missing rather than measured values, which is taken into account where these variables are used to define clamp percentiles. Each behavioral variable was then z-scored using per-variable statistics (mean, standard deviation) computed separately for each recording session.

### Digital twin model architecture

We developed a neural network—the Behavioral Twin—that jointly predicts V1 spike counts from the visual stimulus and concurrent behavioral state. The model receives as input a short temporal clip of grayscale video frames (sequence length *T* = 5 frames, 60 × 80 pixels, ~240 ms) from the head-mounted world camera and a corresponding sequence of six z-scored behavioral variables.

#### Gaze-contingent spatial transformer (Shifter)

To account for gaze-dependent receptive-field shifts, the model includes a learned spatial transformer module, following the approach of Walker et al. (2019); Parker et al. (2022); Xu et al. (2023). The Shifter takes four gaze signals per frame (*θ, ϕ*, pitch, roll) and passes them through a three-layer fully connected network (hidden dimension 256, batch normalization and tanh activations at each layer), outputting per-frame horizontal translation, vertical translation, and rotation parameters. A learnable bias is added before scaling (horizontal: ×80*/*5.5 pixels, vertical: ×60*/*5.5 pixels, rotation: ×180*/*4 degrees). These parameters are used to construct an affine transformation matrix via the Kornia library (Riba et al., 2020), which warps each input video frame to a gaze-corrected reference frame. The Shifter is trained end-to-end with the rest of the model.

#### 3D convolutional visual encoder

The visual feature extractor is a 3D convolutional network with causal temporal structure, built from DenseNet-style blocks. It consists of:

1. A convolutional stem: Conv3D(1 → 48, kernel 3 × 7 × 7, stride 1 × 2 × 2) followed by GELU activation.
2. Three causal dense blocks, each contributing 24 growth channels. Each block uses a bottleneck architecture: (i) a 1 × 1 × 1 pointwise convolution expanding to 4 growth channels, (ii) a temporal convolution (kernel 3 × 1 × 1) with causal zero-padding (*k* −1 frames prepended, zero future padding), and (iii) a spatial convolution (kernel 1 × 7 × 7) with symmetric padding. All convolutions are followed by batch normalization and GELU activations. Dropout (*p* = 0.3) is applied after each block. Spatial max-pooling (1×2×2) is applied between blocks.
3. A behavior-modulated non-local attention mechanism, inserted between the second and third dense blocks (see below).

After the final dense block and pooling, spatial dimensions are collapsed via global average pooling, yielding a temporal sequence of 120-dimensional feature vectors (48 initial + 3 × 24 growth channels).

#### Behavior-modulated non-local attention

Between the second and third encoder blocks, a split spatiotemporal non-local attention module computes self-attention separately along temporal and spatial dimensions. Critically, the query and key representations are conditioned on the behavioral state via Feature-wise Linear Modulation (FiLM): learned linear projections of the time-averaged behavioral vector produce per-channel gain (*γ*) and bias (*β*) terms that multiplicatively and additively modulate the query and key maps before the attention computation. This is applied independently to both the temporal and spatial attention pathways. A learned sigmoid gate, also derived from the behavioral state, regulates the overall strength of the attention residual via the operation 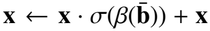 where 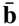 is the mean behavioral state over the sequence. This mechanism allows behavioral context to reshape the model’s internal visual representations at an intermediate processing stage.

#### Behavioral fusion and readout

Visual features are projected to a 512-dimensional space via a linear layer. The six-dimensional behavioral state vector is concatenated with the visual feature sequence and processed by a recurrent (LSTM) module (hidden dimension 256). The final hidden state is passed through a mouse-specific linear readout layer followed by a softplus nonlinearity, producing predicted firing rates for all recorded neurons of the given mouse.

#### Training procedure

The model was trained to minimize the Poisson negative log-likelihood loss between predicted and observed spike counts:

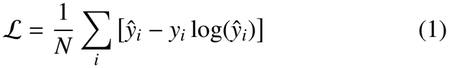

where *ŷ*_*i*_ is the predicted firing rate (after softplus) and *y*_*i*_ is the observed spike count.

An L1 regularization penalty was applied to the mouse-specific readout weights to encourage sparse neural readouts (*λ*_L1_ = 1.0, normalized by the number of readout parameters). Training used the Adam optimizer (Kingma and Ba, 2015) with a maximum learning rate of 4 × 10^−4^, following a OneCycleLR schedule over the full training run. Mini-batch size was 256 samples. Mixed-precision training (float16 via PyTorch AMP) was used to accelerate computation, with gradient scaling. Gradients were clipped to a maximum norm of 5.0.

At each training step, a mouse was selected uniformly at random from the three recording sessions, and a single batch was drawn from that mouse’s training set. This random interleaving ensured balanced exposure across mice without explicit over-sampling. Training proceeded for up to 100 epochs with early stopping (patience = 10 epochs without improvement in validation loss). The model checkpoint with the lowest validation loss was retained for all subsequent analyses.

Input video frames were normalized to [0, 1] (per-segment min–max normalization). The model predicts the spike count at one frame ahead of the final input frame (predict_offset = 1), corresponding to a ~ 48 ms prediction horizon.

Model performance was assessed by the zero-lag cross-correlation (cc) between predicted and observed spike counts on the held-out test set. Correlations were computed both on raw time series and on smoothed time series (boxcar filter, 2 s window, ~42 frames).

#### *Retrained models and shu*ffl*ed controls*

To test whether the behavioral effects depend on one particular training run, ten additional models were trained from scratch with identical architecture, data, preprocessing and hyperparameters, differing only in the random seed. All models in this set were replayed on a coarser stimulus grid than the canonical replay: 36 orientations in 10^°^ steps, the same spatial-frequency set, and two contrasts (0.714 and the maximum). All ten met the same early-stopping criterion as the model used throughout, and reached comparable prediction accuracy (pooled Pearson correlation 0.311 ± 0.002 across seeds, range 0.306 to 0.315, against 0.309 for the model shown in the main figures). The spread across these models therefore reflects run-to-run variability of the training procedure as well as the choice of seed, which is the relevant null for asking whether an effect follows from the recordings rather than from one optimization. Three further control models were trained identically except that the locomotion trace was shuffled in time, breaking its relationship to the video and the neural activity while leaving the architecture, the other behavioral variables and the training procedure intact; in these controls the locomotion gain falls to between −0.45 and − 0.02 Hz and the high-spatial-frequency bias to between +0.07 and +0.10. All models were replayed with the same clamps and analyzed identically.

### In silico experiments

To characterize the visual tuning properties learned by the digital twin and to assess how behavioral state modulates these properties, we performed a series of *in silico* experiments. The trained model was frozen, and synthetic drifting gratings were presented while behavioral variables were clamped to specific values.

#### Stimulus generation

Drifting sinusoidal gratings were generated as sequences of 60 × 80 pixel frames at the training frame rate (dt = 48 ms). Each trial consisted of a blank period (5 frames at mean luminance 0.5) followed by a grating period (70 frames). Pixel luminance was computed as:

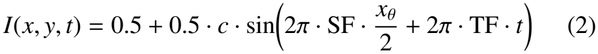

where *c* is contrast, SF is spatial frequency (in cycles per image width), *x*_*θ*_ is the spatial coordinate rotated by the grating orientation, and TF is temporal frequency. Gratings were parameterized by:

- Orientations: 72 values from 0 to 355^°^ in 5^°^ steps.
- Spatial frequencies: 11 values in cycles per image (1, 2, 3, 4, 6, 8, 12, 16, 20, 24, 32), converted to cycles per degree by dividing by 120 (the 80-pixel image width subtends 120^°^). For the analysis of locomotion gain as a function of spatial frequency (Figure 4H), the grid was extended by two further values (28 and 38.4 cycles per image) to resolve the peak of the gain curve; the displayed axis is truncated at 32 cycles per image (0.267 cycles per degree).
- Contrasts: 8 levels evenly spaced from 0 to 1.
- Temporal frequency: 2 Hz.

The Shifter module was disabled during *in silico* experiments (eye and head-position inputs zeroed), ensuring that grating stimuli were presented in a fixed retinotopic frame.

#### Behavioral clamping

For each experiment, all six behavioral variables were held constant across the full trial duration, and one variable was swept at a time while the others were held fixed, so that any change in the response is attributable to that variable alone.

Clamping values were chosen so that every variable is displaced by the same amount in z-scored units, making the six directly comparable: each was set to its own 10th percentile and then to a value 1.5 standard deviations higher, computed on the distribution pooled across the three recordings. Non-swept variables were held at their mean, except locomotion speed, which was held at 0 cm/s to isolate each variable of interest from locomotion-related confounds.

#### Model inference

For each stimulus condition (orientation × spatial frequency × contrast) and behavioral clamp, the full trial video (75 frames) was processed using a sliding window of *T* = 5 frames, yielding 71 model output time points per trial. Firing rates during the blank and grating periods were averaged separately, and evoked responses were computed as the difference (grating − blank). This procedure was repeated independently for each of the three mice.

### Tuning analysis

#### Neuron inclusion

Neurons were included if their prediction correlation, the cross-correlation cc between predicted and recorded responses on the held-out test set, exceeded 0.15. By this criterion, 212 of 223 neurons were included. The threshold is applied to the unsmoothed correlation and was kept permissive deliberately: prediction accuracy is somewhat higher for narrow-than for broad-waveform neurons (0.34 versus 0.29), so a stricter criterion would enrich the narrow-waveform class and bias the cell-type comparisons. All cell-type differences reported for Figure 2 hold at every threshold between 0 and 0.25.

#### Orientation selectivity

Orientation tuning curves were constructed by averaging evoked firing rates across all spatial frequencies and the two highest contrasts. The orientation selectivity index (OSI) was computed as the magnitude of the normalized circular mean of the response vector:

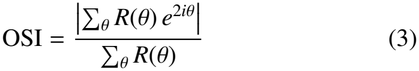

where *R*(*θ*) is the baseline-subtracted response at orientation *θ*. The direction selectivity index (DSI) was computed analogously without the factor of 2 in the complex exponential. Preferred orientation was estimated from the circular mean.

Tuning width was obtained by fitting a double-peaked Gaussian (peaks separated by 180^°^) to the tuning curve via nonlinear least-squares:

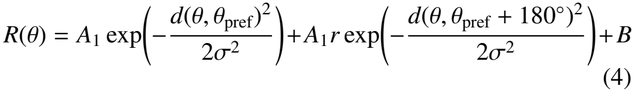

where *d*(·,·) is circular distance in degrees, *r* is the ratio of the secondary to primary peak amplitude, *B* is a constant baseline, and *σ* defines the tuning width. Following Niell and Stryker (2008) and Atallah et al. (2012), tuning width is reported as the half-width at half-maximum of the fitted curve above the baseline, 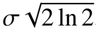 ln 2, so that values are directly comparable with those studies. Fits were run from multiple starting values, and neurons were included if the fit explained at least 70% of the response variance.

#### Spatial frequency selectivit

Spatial frequency tuning curves were constructed at orientations within ±15^°^ of the preferred orientation, at the maximum contrast. A difference-of-Gaussians (DoG) model was fit to each curve:

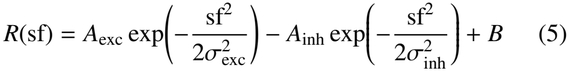

Preferred spatial frequency was defined as the peak of the fitted curve. Bandwidth was computed as the full width at half-maximum in octaves. Neurons were classified as low-pass if the fitted peak occurred at the lowest tested spatial frequency. This peak-position criterion defines the band-pass set used for the cell-type comparisons of Figure 2 and Figure S2 (*n* = 193). The analyses of behavioral modulation in Figure 4 additionally require the full-field-flicker criterion described below, giving a smaller set (*n* = 145). Difference-of-Gaussians fits that failed to converge were excluded; no *R*^2^ threshold was applied.

#### Contrast response

Contrast response functions were extracted at the preferred orientation and spatial frequency. A Naka-Rushton function was fit:

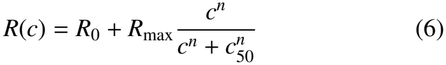

yielding the contrast exponent *n* and semi-saturation contrast *c*_50_. Cell-type comparisons of these tuning metrics used a two-sided permutation test on the difference of group means with 20,000 permutations and an add-one estimator, so the smallest attainable *p* value is 5 × 10^−5^; comparisons of preferred spatial frequency were made on log-transformed values, matching the geometric means reported.

### Behavioral modulation statistics

For each pair of behavioral conditions (e.g., stationary vs. running), within-neuron changes in tuning metrics were assessed using the Wilcoxon signed-rank test (two-sided). Group comparisons (e.g., gain in neurons with low vs. high preferred spatial frequency) used the Mann-Whitney *U* test. All tests were conducted across the population of neurons passing the inclusion criterion. Effect sizes are reported as median differences.

The comparison across behavioral variables (Figure 4I,J) is made at the level of trained networks rather than neurons. For each of the ten retrained digital twins the effect of a variable is summarized as the median across neurons; the ten resulting values are then summarized by their mean with a *t*-based 95% confidence interval and tested against zero with a one-sample *t*-test. This is the appropriate level for that comparison because the question is whether an effect follows from the recordings rather than from one optimization: within a single deterministic network, a signed-rank test over a few hundred neurons detects shifts far smaller than the spread the same variable shows across independently trained networks.

### Behavioral modulation index

To quantify how each behavioral variable modulates V1 firing, the model was probed with each variable clamped at 11 values spanning −2 to +2 standard deviations (dense near zero), with all other variables held at baseline. For each neuron, a modulation index was computed as MI = (*R*_+2_ − *R*_−2_)*/*(|*R*_+2_| + |*R*_−2_ |), where *R*_+2_ and *R*_−2_ are the evoked responses at the high and low clamp; MI lies in [− 1, 1], with positive values indicating that higher values of the variable increased firing.

Significance was assessed by a permutation test in which the clamp positions were shuffled relative to the response curve (1000 permutations). The test statistic was the larger of (i) the absolute Spearman correlation between clamp value and response, capturing monotonic tuning, and (ii) the absolute Spearman correlation between |*z*| and the response deviation from its value at *z* = 0, capturing symmetric, U-shaped tuning. Perneuron *p*-values were corrected across all neurons and variables with the Benjamini–Hochberg procedure (false-discovery rate 0.05).

For each variable, the population summary reported in Figure 3C is the mean of the per-neuron signed MI (whether the population shares a direction of modulation) plotted against the mean of |MI | (how strongly the variable modulates individual neurons), taken across all *N* = 212 neurons.

### Behavioral modulation of visual tuning

To test how behavioral state reshapes visual tuning, drifting gratings were replayed from a common stationary ground state while a single behavioral variable was clamped, and tuning metrics (evoked firing rate, preferred spatial frequency, spatial-frequency bandwidth) were compared between states.

Spatial-frequency analyses were restricted to band-pass neurons: neurons with a converged difference-of-Gaussians fit whose peak fell within the tested range, and whose response to a uniform flickering field was below 65% of the response at the preferred spatial frequency, the latter excluding neurons that are not spatially tuned within the tested range. The band-pass set was defined once from the common ground state and applied to both states being compared (*n* = 145). Locomotion gain as a function of spatial frequency was computed per neuron as the ratio of running to stationary responses at each spatial frequency (maximum contrast, averaged across orientation) and summarized as the population median with its interquartile range.

The locomotion-specific shift toward high spatial frequencies was quantified per neuron as the difference between the locomotion gain above and below that neuron’s own preferred spatial frequency: the mean gain at spatial frequencies above twice the preferred value minus the mean gain below half of it, computed for the *n* = 138 band-pass neurons with responses in both zones. Contrasting the two flanks rather than referencing the peak keeps the compared responses matched in baseline drive, so the statistic reflects spatial frequency rather than overall response level.

### Generalization to the Allen Brain Observatory

The trained model was evaluated on the Allen Brain Observatory Visual Coding dataset (Neuropixels, `brain_observatory_1.1`). Recordings from 26 sessions with at least 30 V1 (VISp) units were used. Units were included on the basis of response reliability rather than response variance: for each unit the split-half correlation of its movie response across repeat presentations was computed over 20 random splits of the trials and Spearman–Brown corrected, and units reaching a corrected reliability of 0.3 were retained (1726 of 1841 units; median reliability 0.71). Sessions were required to retain at least ten such units, which all 26 did. The natural-movie stimulus was resized to 60 × 80 pixels and replayed through the frozen model with all behavioral inputs set to zero, and the predicted population responses were compared with the trial-averaged recorded responses (49 ms bins).

Allen movies were presented to the frozen twin with all behavioral inputs set to zero. Population geometry was compared by representational similarity analysis. For each population, a representational dissimilarity matrix (RDM) was built across 120 subsampled movie time bins using correlation distance (1 − Pearson correlation), and the similarity between two RDMs was the Spearman correlation of their upper triangles. The twin-to-Allen similarity (pooled across the three model mice) was compared with the inter-mouse similarity computed over all 325 pairs of the 26 sessions, with a generalized linear model (GLM) predicting responses from the visual input, trained on the same freely moving data, and with a randomly initialized model as a lower bound.

Because representational similarity is attenuated by RDM unreliability, all similarities were disattenuated before comparison (**?**): each RDM’s split-half reliability was estimated and Spearman–Brown corrected (mean 0.76 across sessions), model-to-animal similarities were divided by the square root of the animal RDM’s reliability, and animal-to-animal similarities by the square root of the product of both reliabilities. This matters because the two comparisons are attenuated by different amounts, so the uncorrected values are not directly comparable.

The twin was compared with the GLM by a paired test across the 26 sessions. Equivalence between the twin and the inter-mouse distribution was assessed by two one-sided tests, with the equivalence margin set to the standard deviation of the mouse-to-mouse difference distribution; the twin’s rank within that distribution is reported as a percentile. The per-session correspondence between twin-to-Allen and inter-mouse similarity was quantified with Spearman correlation. As an independent check that does not rely on the reliability correction, the analysis was repeated with model predictions resampled as Poisson counts at each session’s repeat number and trial-averaged, matching the measurement noise of the recordings.

### Software

All models were implemented in PyTorch. Affine image transformations used the Kornia library (Riba et al., 2020). Tuning curve fits used SciPy’s `curve_fit`. Statistical tests used SciPy. Figures were generated with Matplotlib and Seaborn.

---

**Figure S1:**
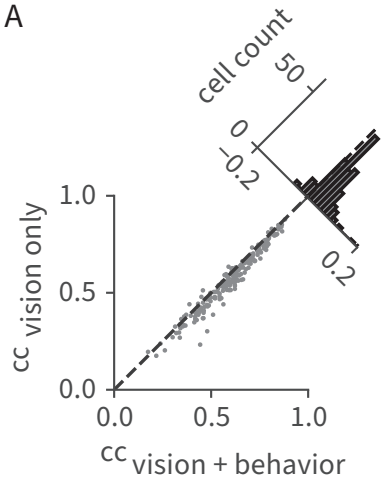
Behavioral inputs improve single-neuron prediction accuracy. Single-neuron prediction accuracy (Pearson correlation between predicted and recorded spike counts on held-out data) for the full visual-behavioral model versus a control in which the behavioral modulation pathway is bypassed, leaving the visual encoder and the gaze-correcting shifter intact. Each point is one neuron; the marginal histogram shows the per-neuron difference (full minus vision-only). Including behavior improved prediction for the large majority of neurons.

**Figure S2:**
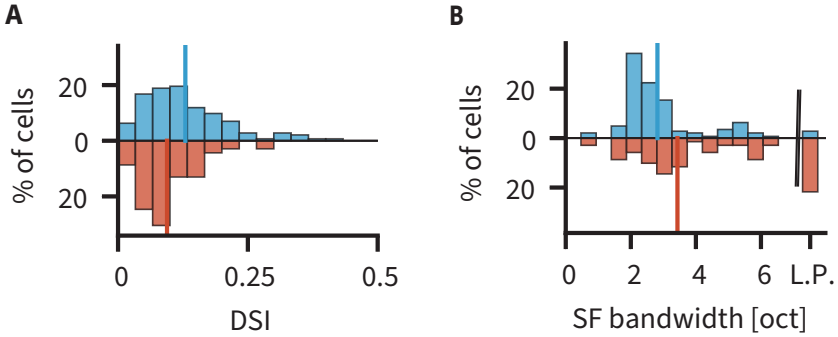
Cell-type differences in direction selectivity and spatial-frequency bandwidth. **(A)** Direction selectivity index for broad-waveform (BW, putative excitatory) and narrow-waveform (NW, putative inhibitory) neurons. BW neurons are more direction selective (mean DSI 0.130, *n* = 143, versus 0.094, *n* = 69; permutation test on the difference of means, *p* = 9 × 10^−4^). **(B)** Spatial-frequency bandwidth for the two cell types, band-pass neurons only. NW neurons are more broadly tuned (mean 3.43 octaves, *n* = 54, versus 2.82, *n* = 139; *p* = 0.003).

**Figure S3:**
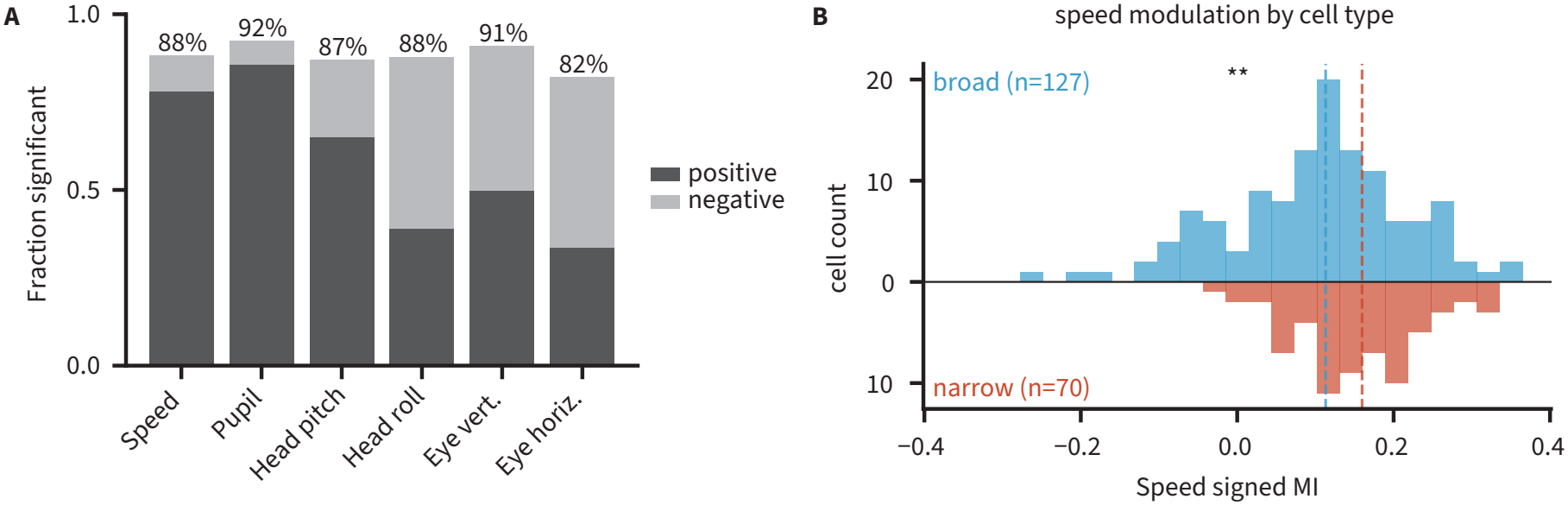
Prevalence and cell-type specificity of behavioral modulation. **(A)** Fraction of neurons significantly modulated by each variable (shape-aware permutation test), split into positively and negatively modulated cells; the percentage above each bar gives the total. Locomotion speed and pupil are predominantly positive, whereas the head- and eye-position variables are mixed. **(B)** Distributions of the per-neuron locomotion-speed modulation index for broad-waveform (BW, putative excitatory) and narrow-waveform (NW, putative inhibitory) neurons. Narrow-waveform neurons are more strongly speed-modulated (median MI 0.17, *n* = 67, versus 0.11, *n* = 123; Mann–Whitney U, *p* = 4.2 × 10^−4^; significantly modulated neurons only). The effect is specific to locomotion speed: the same test applied to each of the other five variables gives *p* ≥ 0.39.

